# Proteus mirabilis entails a switch from commensal to pathogen - Genomic insights from a blaTEM-1B-harboring novel isolate from India

**DOI:** 10.1101/2025.11.25.690331

**Authors:** Sovon Acharya, Parmanand Kushwaha, Shailesh Desai, Soham Biswas, Langamba Angom Longjam, Biswaroop Chatterjee, Munindra Ruwali, Sachinandan De, Lakshminarasimhan Krishnaswamy, Prashanth Suravajhala, Gyaneshwer Chaubey

## Abstract

**Background:** *Proteus mirabilis* is an opportunistic pathogen that can shift from gut commensalism to causing severe urinary tract and bloodstream infections. The genomic basis of *P. mirabilis*’ transition from commensalism to severe infection is not well understood, especially in Indian clinical isolates. This study aimed to explore the genomic attributes of a prevalent *P. mirabilis* strain in India.

**Methods:** Whole-genome sequencing (WGS) was performed on a clinical *P. mirabilis* isolate by using NGS. AMR genes and virulence factors were identified using ResFinder/CARD and VFDB, respectively. Sequence typing was performed with MLST. Protein-protein interactions and noncoding RNAs were analyzed using STRING, CytoHubba, and BAKTA.

**Results:** WGS confirmed the multidrug-resistant nature of the isolate, revealing a broad repertoire of AMR genes, including *blaTEM−1B*, conferring resistance to beta-lactam antibiotics. Several resistance determinants were co-localised with insertion sequences on a single contig, indicating a high potential for horizontal gene transfer. The virulome profile highlighted multiple fimbrial genes, hemolysins, and toxin-encoding loci, together supporting strong adhesion, biofilm formation, and host tissue damage. Comparative analysis revealed unique genomic signatures not present in reference isolates, including additional virulence determinants and mobile genetic elements. MLST profile of the isolate is closely related to ST-675 & ST-797, while virulence typing (vST-1157) indicated a novel profile distinct from reference ST-675 strains. Insilico protein–protein interaction of this isolate revealed a highly interconnected virulence network, with urease, fimbrial adhesins, and hemolysins emerging as central functional modules driving pathogenicity. In addition, a noncoding RNA was identified within the genomic context of the ure operon (Contig 10), suggesting a possible regulatory role in urease expression. PPI network analysis highlighted urease, fimbrial proteins, and hemolysins as central hubs contributing to pathogenicity.

**Conclusions:** This study presents the first genomic characterization of a multidrug-resistant *P. mirabilis* clinical isolate from India, designated as *P. mirabilis* strain Indica. The convergence of AMR genes, virulence factors, MGEs and a novel noncoding RNAs linked to urease expression highlights the emergence of a high-risk variant in India. These findings have significant implications for infection control and emphasize the need for genomic surveillance to guide therapeutic strategies and develop new interventions globally.

## 1. Introduction

*P. mirabilis* is a Gram-negative bacterium of the family Enterobacteriaceae, widely recognized for its dual identity as both a harmless commensal of the human gut and a significant opportunistic pathogen. While its presence in the gut microbiota is often benign, under certain conditions, it can breach anatomical barriers and cause a range of severe infections in humans. Among these, urinary tract infections (UTIs) are its most well-known manifestation, particularly complicated cases and those associated with long-term indwelling catheters, known as catheter-associated UTIs (CAUTIs) (1–3). The bacterium’s ability to thrive in the urinary tract is a consequence of its unique virulence traits, which facilitate colonization, persistence and tissue damage. Beyond UTIs, *P. mirabilis* has been implicated in more serious systemic infections, including wound infections, bacteremia and septicemia, which can be life-threatening, especially in vulnerable patient populations such as the elderly, diabetics, cancer patients and immunocompromised individuals (4, 5). Understanding the genomic basis of its transition from a commensal to a virulent pathogen is crucial for developing effective therapeutic and preventative strategies.

The pathogenic success of *P. mirabilis* is rooted in its sophisticated and coordinated expression of multiple antibiotic resistance (ABR) genes, virulence factors and mobile genetic elements (MGEs). Its most distinguishing feature is the production of urease, an enzyme that hydrolyzes urea into ammonia and carbon dioxide (6). This enzymatic activity significantly raises the pH of urine, creating a hostile alkaline environment for other bacteria while promoting the precipitation of magnesium and calcium ions (7). This process leads to the formation of crystalline biofilms and struvite (magnesium ammonium phosphate) stones, which can cause severe kidney damage, block urinary catheters and render antibiotic treatment ineffective (8, 9). This unique mechanism is a primary reason for the persistent and recurrent nature of *P. mirabilis* UTIs. Another hallmark of this bacterium is its characteristic swarming motility, a collective form of cell movement across solid surfaces (10). This process involves a reversible differentiation of short, flagellated vegetative cells into elongated, hyperflagellated “swarmer” cells. This swarming behavior allows *P. mirabilis* to rapidly ascend the urinary tract, a critical step in the pathogenesis of pyelonephritis (kidney infection) (11). Furthermore, swarming is intricately linked to biofilm formation, a complex process where bacteria adhere to surfaces, such as urinary catheters and encase themselves in a protective matrix (12). This biofilm matrix provides a physical barrier against host immune defenses and antibiotics, contributing to chronic infections. The ability of *P. mirabilis* to adhere to host cells is mediated by a diverse repertoire of fimbrial adhesins. These include the mannose-resistant *Proteus* (MR/P) fimbriae and the *P. mirabilis* fimbriae (PMF), which bind to specific receptors on uroepithelial cells, facilitating colonization and preventing the bacteria from being flushed out by urine flow (13). Other important virulence factors include hemolysins such as HpmA, which damage host cell membranes and release essential nutrients like iron (14), and proteases, which can degrade host tissues and evade the immune system. Additionally, the bacterium’s robust iron acquisition systems are essential for its survival in the iron-limited environment of the urinary tract, allowing it to scavenge iron from host proteins like transferrin and lactoferrin (15). The collective and synergistic action of these virulence factors allows *P. mirabilis* to successfully colonize, invade and cause severe disease when host defenses are compromised.

Historically, *P. mirabilis* was considered more susceptible to antibiotics compared to other Enterobacteriaceae like *Escherichia coli* or *Klebsiella pneumoniae*. However, over the past few decades, it has rapidly emerged as a significant reservoir of antimicrobial resistance (AMR) (16). This has transformed it from a manageable pathogen into a multidrug-resistant (MDR) threat, severely complicating therapeutic options. A major concern is the widespread prevalence of β-lactamases, enzymes that break down the β-lactam ring of antibiotics such as penicillins and cephalosporins. These include TEM- and CTX-M-type extended-spectrum β-lactamases (ESBLs), which have been globally disseminated, leading to treatment failures (17, 18). While ESBLs have received considerable attention, the growing threat of carbapenemases (e.g., NDM, KPC) in Enterobacteriaceae is also a grave concern, with some rare reports of their presence in *Proteus* species further highlighting the escalating crisis (19). Beyond β-lactamases, *P. mirabilis* has acquired resistance to multiple other antibiotic classes. Genes encoding aminoglycoside-modifying enzymes (e.g., *aph* genes), sulfonamide resistance genes (*sul1*, *sul2*), and tetracycline efflux pumps (*tet* genes) are frequently reported (9). The emergence of plasmid-mediated quinolone resistance (*qnr* variants) is also a growing issue, limiting the use of a key class of antibiotics for treating UTIs (20). The coexistence of these resistance determinants on MGE allows a single isolate to be resistant to multiple drugs, a hallmark of MDR strains. This rapid acquisition of resistance traits has been a major driver of the increasing morbidity and mortality associated with *P. mirabilis* infections.

The swift evolution of *P. mirabilis* from a commensal to a MDR pathogen is largely driven by horizontal gene transfer (HGT), a process that allows bacteria to acquire new genetic material from other organisms or *vice versa*(21). HGT is mediated by MGEs such as plasmids, transposons, and bacteriophages. Plasmids are extrachromosomal DNA molecules that can carry both virulence and resistance genes, enabling their rapid transfer between different bacterial species. The acquisition of large plasmids can fundamentally change a bacterium’s phenotype in a single event, making it both more virulent and more drug-resistant. Genomic islands, often referred to as pathogenicity islands (PAIs), are another key mechanism. These are clusters of genes, often with virulence-related functions, that are integrated into the host chromosome and show evidence of HGT. Comparative genomic studies have revealed that pathogenic strains of *P. mirabilis* contain large PAIs that are absent in their commensal counterparts (13, 22). The co-localization of resistance genes and virulence factors on these MGEs underscores a key evolutionary dynamic: the traits that enable a bacterium to survive in a hospital environment (AMR) are often acquired alongside the traits that allow it to cause disease (virulence), leading to the co-evolution of these two critical phenotypes.

Despite the global recognition of *P. mirabilis* as a clinically relevant pathogen, there is a significant lack of comprehensive genomic data from India (Fig.6). The country is grappling with an immense burden of MDR Gram-negative infections, driven by factors such as high population density, widespread use of antibiotics, and limited infection control measures (23). While surveillance efforts in India have predominantly focused on ESBL and carbapenemase-producing *E. coli* and *K. pneumoniae*, the rising clinical significance of *P. mirabilis* warrants urgent genomic characterization (24, 25). Very few studies were performed on whole-genome sequencing (WGS) of Indian *P. mirabilis* isolates, leaving a critical gap in our understanding of their resistome, virulome and evolutionary dynamics. In this study, we aimed to bridge this knowledge gap by providing a detailed whole-genome characterization of a blaTEM−1B–harboring *P. mirabilis* isolate from India. Through the integration of genomic approaches encompassing resistome and virulome profiling, MGE characterization and comparative genomics, this study provides new insights into the genetic mechanisms underlying the evolution of this species from a benign commensal to a multidrug-resistant and highly virulent pathogen. Our findings highlight the critical role of genomic surveillance in monitoring the emergence of high-risk clones and emphasize its value in guiding more effective strategies for diagnostics, therapeutic interventions and infection control both in India and globally.

## 2. Methods

### 2.1. Genome assembly, annotation, and bioinformatic analysis

Raw sequencing reads were first assessed for quality using FastQC (https://www.bioinformatics.babraham.ac.uk/projects/, last accessed May 15, 2025) (14). Adapter trimming and removal of low-quality bases were performed with FastP (15). High-quality reads were subsequently assembled with Unicycler (17) v.0.8.4.0 (https://github.com/rrwick/Unicycler, last accessed Aug 15, 2025), and assembly quality was evaluated using Quality Assessment Tool for Genome Assemblies (QUAST) (18). Genome annotation was initially carried out with Prokka (19) and cross-validated with the Rapid Annotation using Subsystem Technology (RAST) server (33) (https://rast.nmpdr.org/, last accessed Aug 15, 2025). The sequence is deposited at BioProject ID: PRJNA1206681.

For typing and resistance profiling, multilocus sequence typing (MLST) was performed using the PubMLST (34) database (https://pubmlst.org/, last accessed Aug 15, 2025). Antimicrobial resistance genes were identified using tools from the Center for Genomic Epidemiology (http://www.genomicepidemiology.org/, last accessed Aug 15, 2025), including ResFinder and the Comprehensive Antibiotic Resistance Database (CARD) (23, 24). Additional analyses were carried out on the Bacterial and Viral Bioinformatics Resource Center (BV-BRC) platform (https://www.bv-brc.org/, last accessed Aug 15, 2025), which provided validation and complementary insights (35).

The MGEs were detected using ISEScan (22). IslandViewer4 (21) facilitated the identification and visualization of genomic islands. Virulence-associated genes were identified through the Virulence Factor Database (VFDB) (https://ngdc.cncb.ac.cn/databasecommons/database/id/516, last accessed Aug 15, 2025) (25,27). Pairwise sequence comparisons were conducted using NCBI-BLAST, while AlphaFold was employed for protein structure prediction (36).

Raw sequence data quality was assessed with *FastQC* and trimmed with *FastP*. De novo assembly was performed using *Unicycler* and evaluated with *QUAST*. The draft genome was annotated with *Prokka*. Multilocus sequence typing (MLST) was conducted using the PubMLST database (34). While the AMR genes were identified using *ResFinder* and validated with the CARD database, virulence-associated genes were screened against the *Virulence Factor Database* (VFDB). The final list of genes were assessed for pathways using the STRING database (37) and the top ranking genes were visualized and deciphered using Cytoscape-Cytohubba (38).

### 2.2. Comparative Genomics and Phylogenetic Analysis

To understand the evolutionary context, comparative genomics was performed against reference uropathogenic and environmental *P. mirabilis* genomes. Average nucleotide identity (ANI) was calculated to assess genetic relatedness through the autoMLST 2.0 (31). Core genome-based phylogeny was constructed using concatenated alignments of orthologous genes and visualized using iTOL (27). Bioinformatics workflow for WGS analysis was depicted in the figure 1a.

**Figure 1a.**
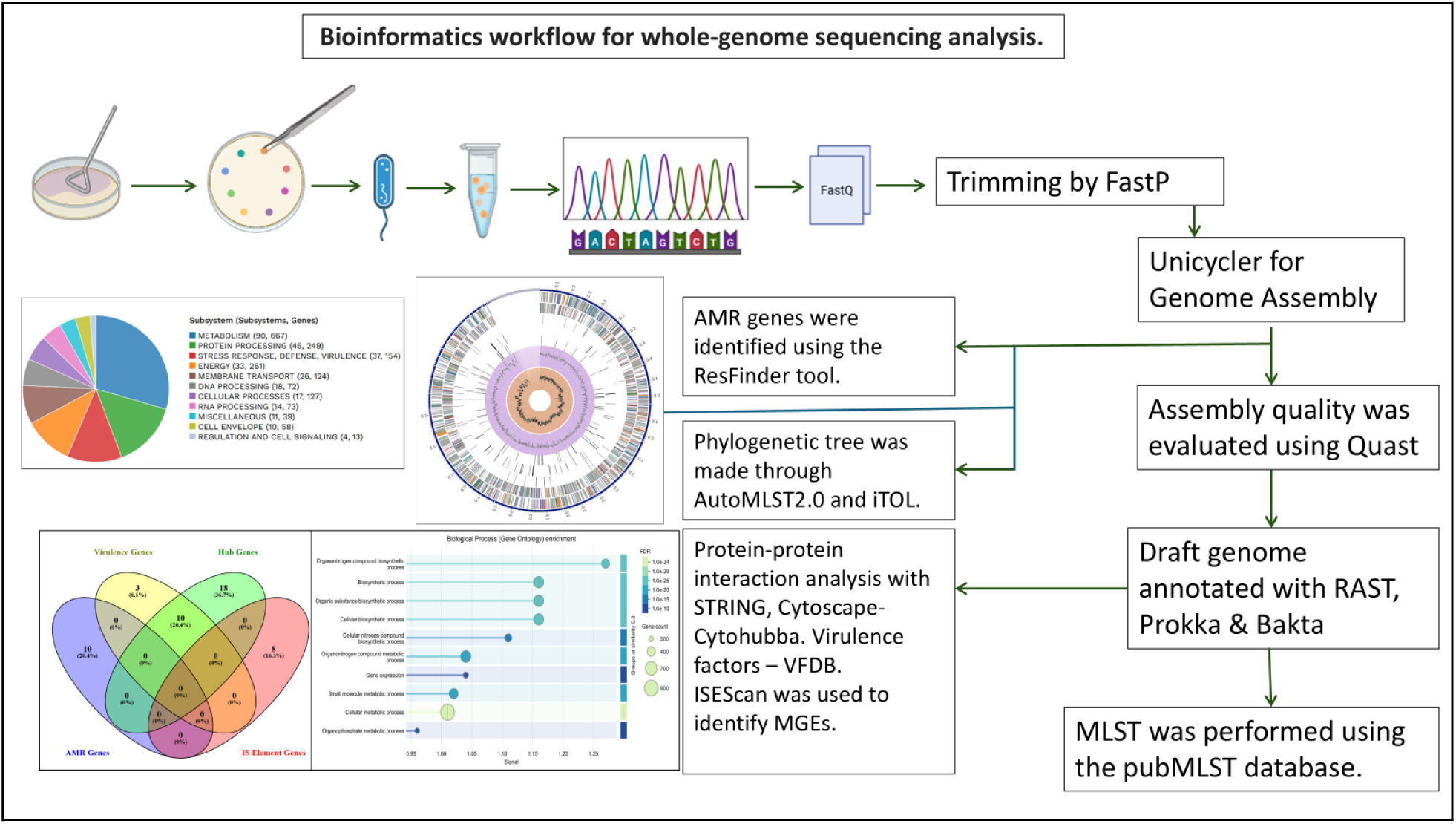
Bioinformatics workflow for whole-genome sequencing analysis.

## 3. Results

### 3.1. Isolation, Identification, and Genomic Characterization led to annotation, yielding a Draft Genome of ∼ 3.9 Mbp and 3652 predicted proteins

Illumina NovaSeq sequencing generated approximately 19.1 million high-quality paired-end reads. The WGS of the *P. mirabilis* isolate yielded a draft genome of approximately 3.9 Mbp with a GC content of 40.5%, assembled into a single circular contig from 19.1 million paired-end reads. While annotation with Prokka revealed 3,655 predicted genes, including 3,652 protein-coding sequences and 82 RNA genes (5 rRNAs and 79 tRNAs), a circular map of the genome (Fig. 1b) provided a visual overview of these features, including GC content, GC skew and the location of predicted genes.

**Figure 1b.**
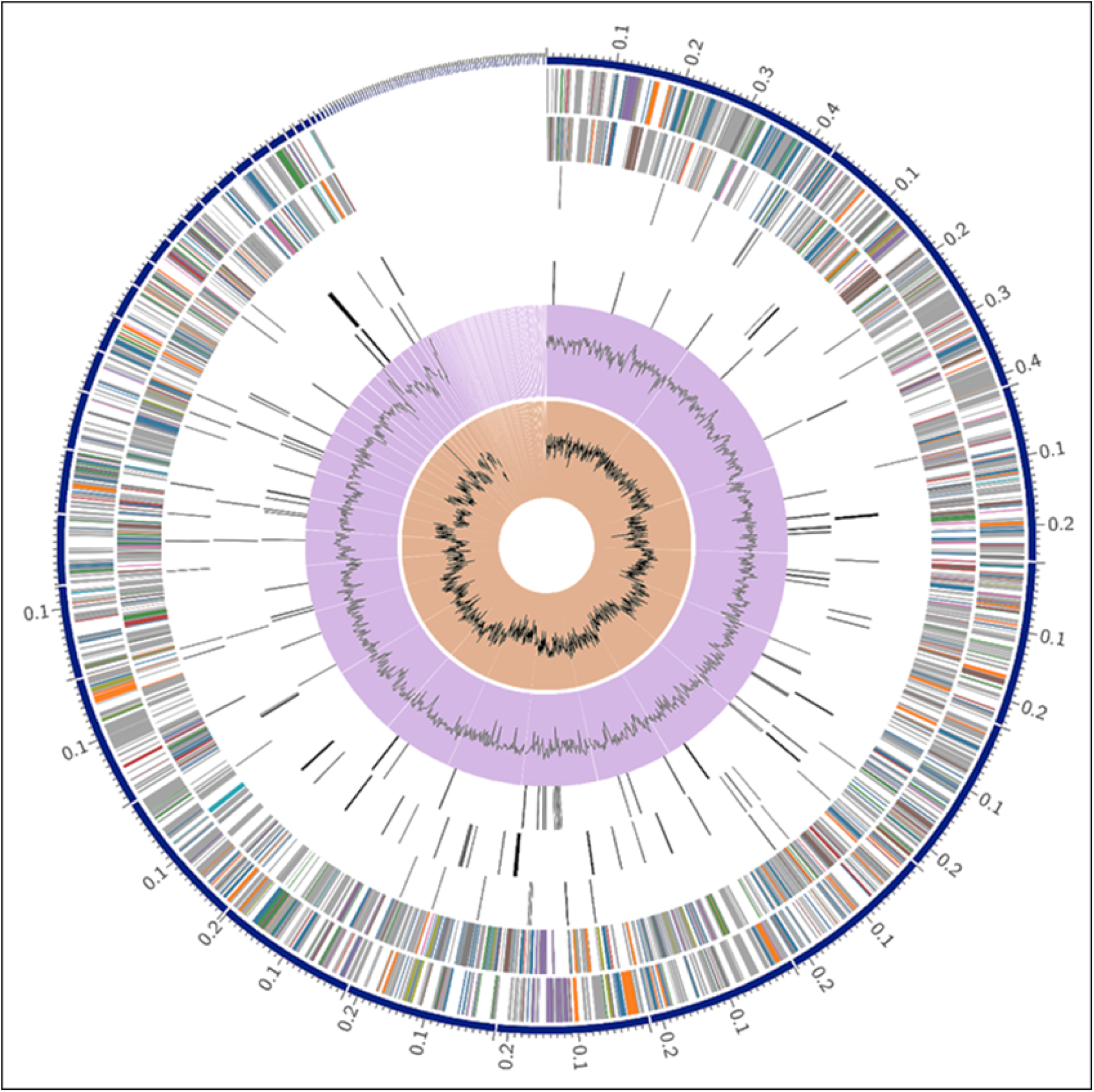
Circular Genome Map of the *P. mirabilis* Isolate.

A circular graphical display of the genome annotations for the *P. mirabilis*. From the outermost to the innermost rings, the figure displays: contigs, coding sequences (CDS) on the forward and reverse strands (colored by subsystem), RNA genes, CDS with homology to known AMR and virulence factors, GC content and GC skew.

### 3.2. Distinct Functional Classification and Subsystems are Associated with *P. mirabilis*

To understand the functional profile of our *P. mirabilis* isolate, we classified its genes into subsystems using the RAST annotation system. The resulting pie chart (Figure 2) shows the distribution of gene functions across key biological categories. Other key functional categories, such as Energy, Membrane Transport and DNA Processing are also well-represented, collectively painting a comprehensive picture of the isolate’s genetic blueprint. This functional breakdown provides context for the virulence and resistance mechanisms identified through other analyses, suggesting that these traits are deeply integrated into the organism’s core biology.

**Figure 2a.**
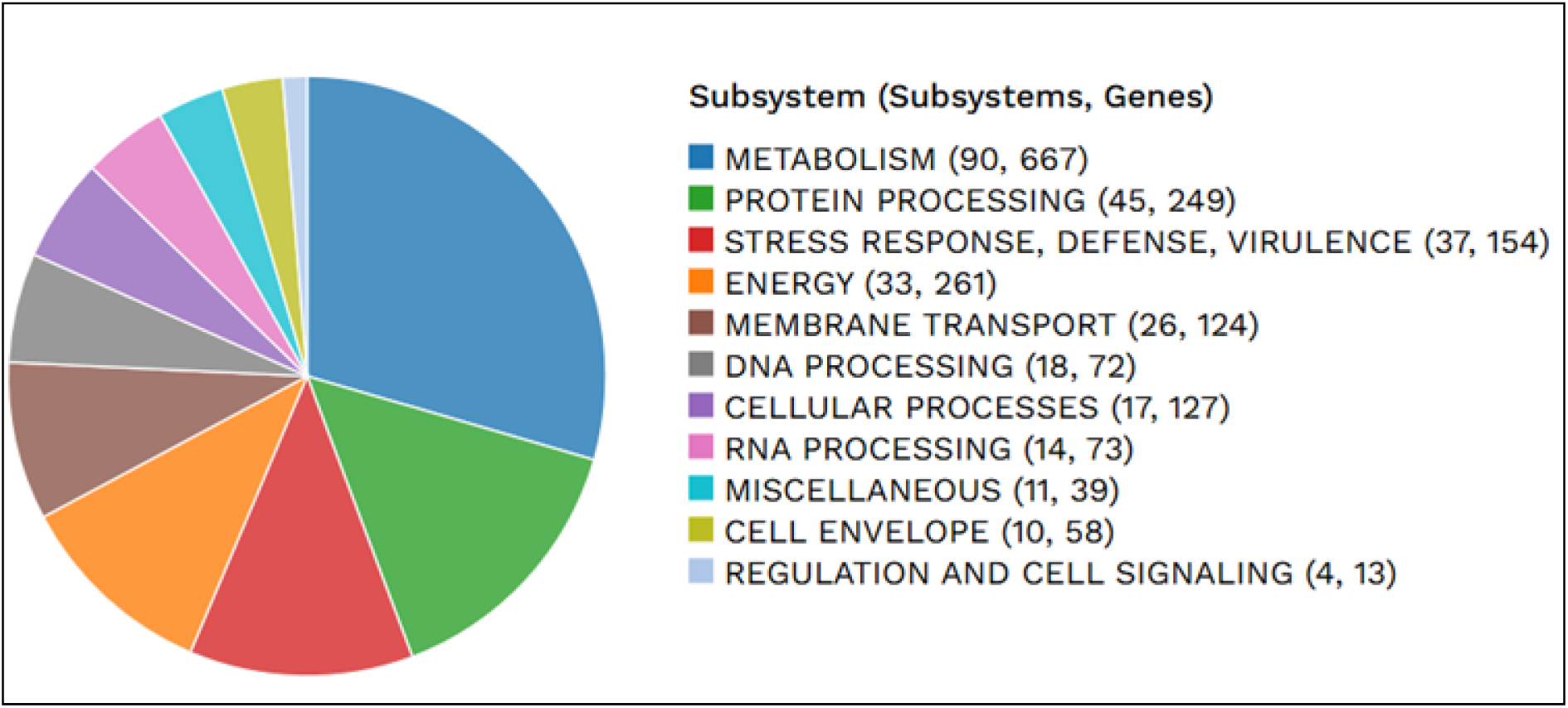
Functional Classification of the *P. mirabilis* Genome. A pie chart illustrating the distribution of the isolate’s genes into functional subsystems. The chart highlights the major functional categories, including Metabolism, Protein Processing, and Stress Response, Defense and Virulence. The number of subsystems and corresponding genes are provided for each category.

Functional enrichment analysis (Fig. 2b)of the *P. mirabilis Indica* genome using STRING revealed significant overrepresentation of biosynthetic and metabolic processes, particularly those involving organonitrogen compounds and cellular biosynthesis. These functions align with the well-established role of *P. mirabilis* in nitrogen metabolism, notably through urease activity that facilitates urinary tract colonization and stone formation. The enrichment of diverse metabolic pathways, including small-molecule and organophosphate metabolism, underscores the organism’s metabolic versatility, enabling persistence in both hospital environments and host tissues. Notably, enrichment in gene expression machinery suggests a genomic bias toward rapid adaptation and response to environmental stressors. Collectively, these enriched biological processes reflect the dual strategy of *P. mirabilis*: sustaining robust metabolic fitness while leveraging specialized nitrogen-based pathways to enhance pathogenicity in vulnerable hosts.

**Figure 2b.**
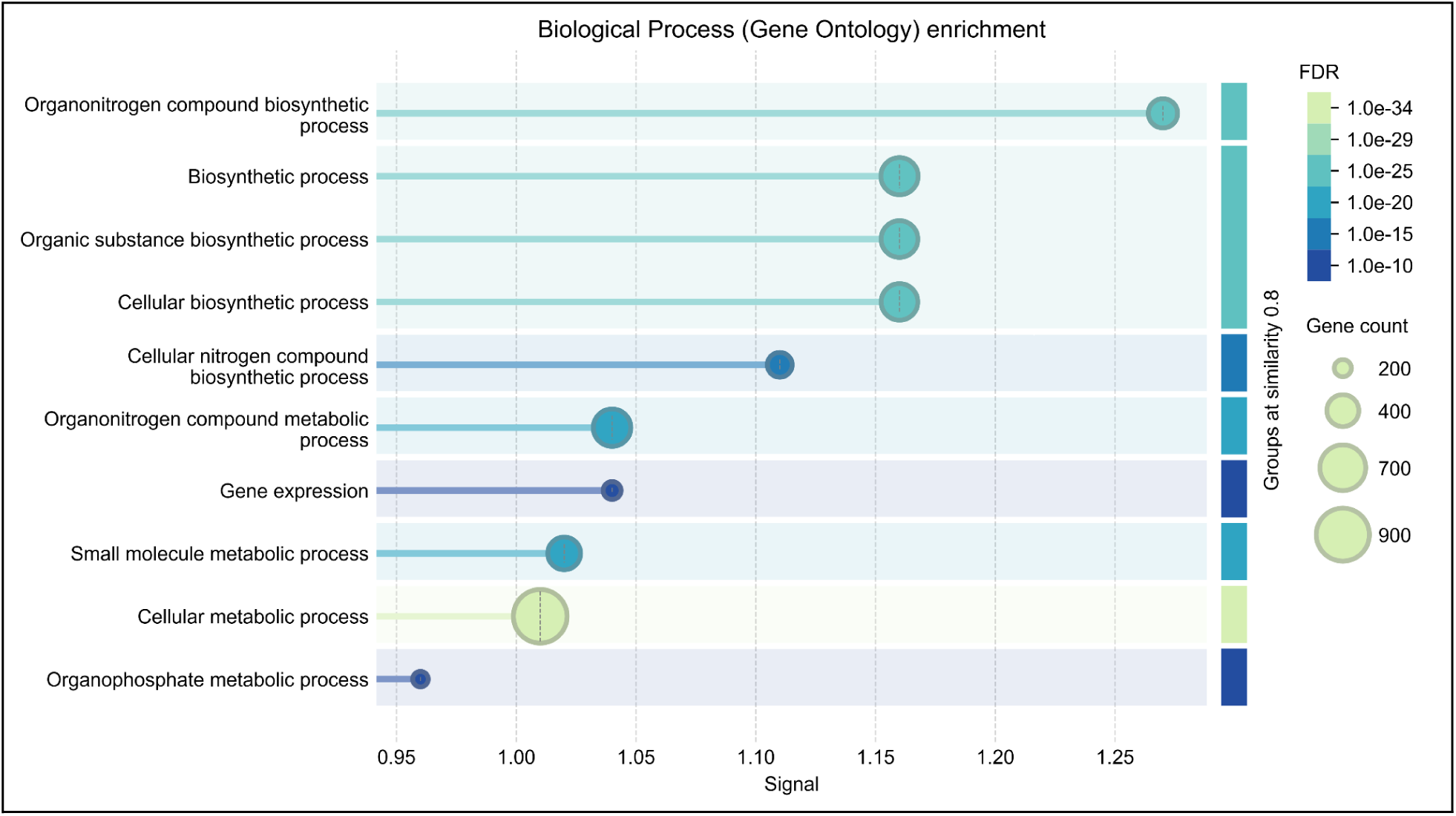
Functional enrichment analysis of *P. mirabilis Indica* isolate.

Biological process Gene Ontology (GO) terms enriched in the genome were identified using STRING. The x-axis represents the enrichment signal, while the y-axis lists significantly enriched biological processes. Bubble size corresponds to the number of genes mapped to each GO category, and bubble color reflects the adjusted *p*-value (FDR), with darker shades indicating stronger significance. Processes related to biosynthesis, organonitrogen metabolism and cellular metabolism were prominently enriched, highlighting the metabolic versatility and nitrogen-driven adaptation strategies of the isolate.

### 3.3. Multilocus Sequence Typing (MLST) ST-675 is attributed to AMR pathogenesis

Taxonomic classification of the assembled genome confirmed the isolate as *P. mirabilis* with 100% confidence. The phylogenetic lineage was resolved as *Pseudomonadota > Gammaproteobacteria > Enterobacterales > Morganellaceae > Proteus > P. mirabilis*. This precise species-level assignment underscores the reliability of the sequencing and classification pipeline and firmly establishes the identity of the isolate as *P. mirabilis*, an opportunistic pathogen frequently implicated in urinary tract infections, catheter-associated infections, and multidrug resistance. Multilocus sequence typing (MLST) analysis assigned the isolate to sequence type ST-675, with five of the six loci (83.3%) matching this profile and a single mismatch placing it in close proximity to ST-797. This phylogenetic positioning suggests that the isolate belongs to the ST-675 lineage but may represent a divergent sub-variant, reflecting ongoing microevolution or horizontal gene exchange within this clonal complex. Virulence typing further revealed a unique profile (vST-1157), characterized by a distinct repertoire of virulence determinants not commonly observed in reference ST-675 isolates. The presence of these additional virulence genes, in combination with AMR loci, underscores the pathogenic versatility of this strain, and is named *P. mirabilis Indica*. Genomic features highlight the dual capacity of the isolate to persist in clinical environments and to cause severe infections, particularly in cancer patients and other immunocompromised hosts, who are highly vulnerable to opportunistic pathogens with both resistance and virulence attributes. Collectively, these findings point to the emergence of a potentially high-risk clonal variant of *P. mirabilis*, linking multidrug resistance with an expanded virulence arsenal and raising concerns for global dissemination and therapeutic challenges. We named this strain P. mirabilis Indica.

### 3.4. The Resistome Blueprint of this Indian isolate shows a substantial difference compared to the isolate collected around 40 years ago in USA

The WGS analysis of the *P. mirabilis* isolate revealed a rich repertoire of acquired AMR genes, which collectively provide a robust genomic basis for its MDR phenotype. This was determined through a comprehensive screening using both the ResFinder and CARD databases. The identified genes are associated with resistance to several clinically important antibiotic classes, highlighting the challenge this isolate poses in a healthcare setting. This analysis revealed the presence of the *blaTEM-1B* gene, which encodes a beta-lactamase enzyme that confers resistance to a wide range of beta-lactam antibiotics, including penicillins (such as amoxicillin and ampicillin) and first- and second-generation cephalosporins. Given its global prevalence, the identification of this gene is a critical indicator of this isolate’s capacity to inactivate common clinical antibiotics and evade treatment. Beyond beta-lactam resistance, the analysis also identified determinants for several other antibiotic classes. Genes such as aph(6)-Id and aph(3′)-Ib, which encode aminoglycoside-modifying enzymes, were identified. These enzymes inactivate streptomycin by preventing its binding to ribosomal targets, thereby rendering the drug ineffective. Resistance to sulfonamides was mediated by the sul1 and sul2 genes, which provide an alternative, drug-insensitive pathway for folic acid synthesis. Furthermore, the cat gene, responsible for chloramphenicol resistance via drug acetylation was also detected. Overall, the genome harbors determinants conferring resistance to multiple antibiotic classes, including two major groups widely employed in the treatment of urinary tract infections.The *tet(J)* gene, encoding an efflux pump that expels tetracycline and doxycycline from the bacterial cell, was found. Similarly, the qnrD3, a plasmid-mediated quinolone resistance determinant, was identified which protects DNA gyrase—the target of fluoroquinolones like ciprofloxacin from inhibition, allowing the bacterium to continue DNA replication even when the drug is present. While the qacE efflux pump gene, which provides resistance to disinfectants was detected, this shows the isolate’s ability to adapt and survive in hospital settings despite common antiseptics. The presence of these resistance genes confirms that this clinical isolate is a multidrug-resistant strain, making treatment more complex (Figure 3 and Table 1). During the annotation of the 1982 *P. mirabilis* isolate (HI4320) genome, only two antibiotic resistance genes were detected: cat and tet. Compared to the resistome profile of the isolate (P. mirabilis Indica) from this study, the genome of HI4320 contained fewer antibiotic resistance genes. This observation aligns with common patterns of antibiotic resistance development and spread in bacterial populations. Typically, earlier isolates tend to have fewer resistance genes, while more recent strains acquire additional resistance genes through horizontal gene transfer over time.

**Figure 3.**
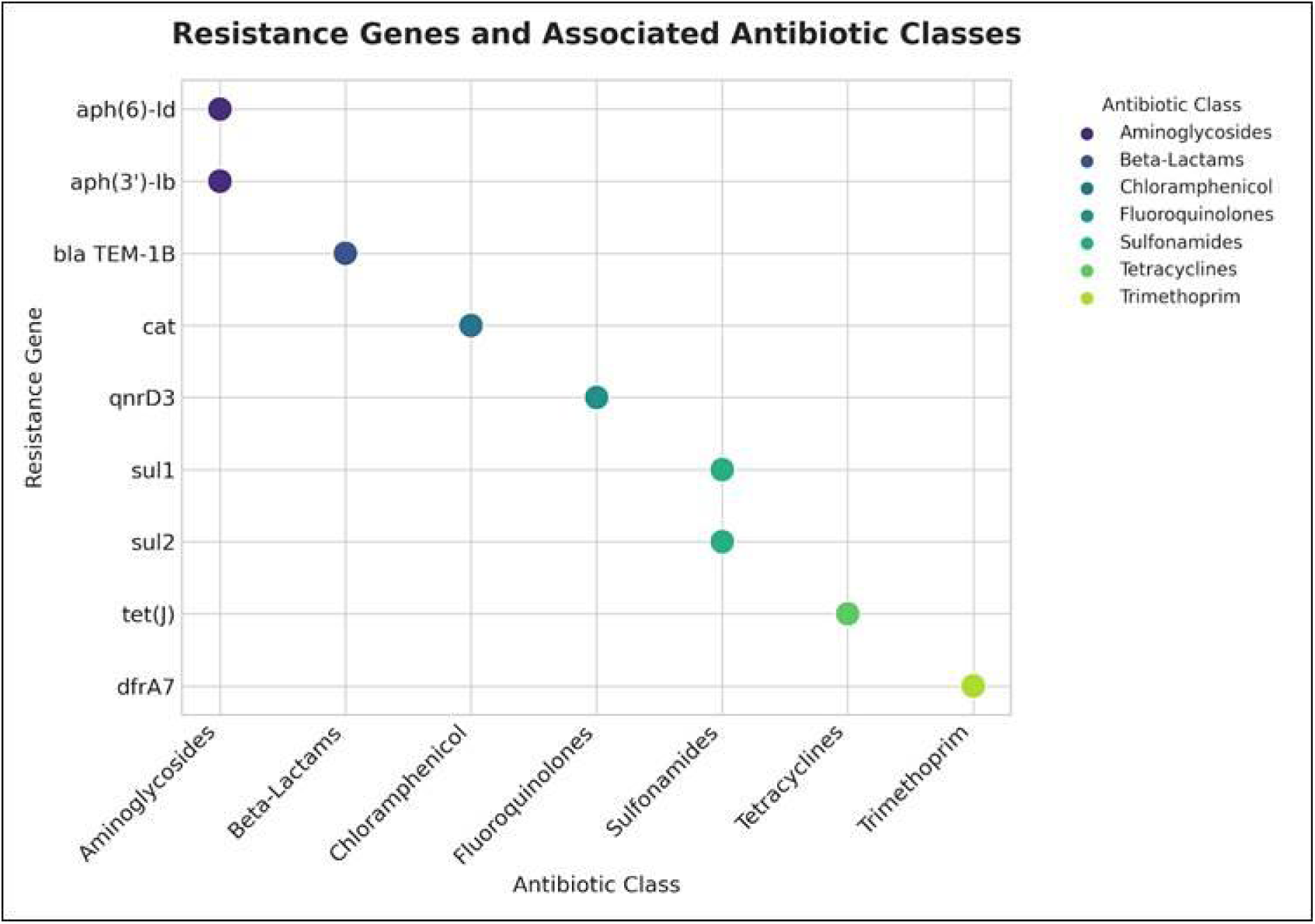
Antibiotic Resistance Gene Profile of the *P. mirabilis* Isolate.

**Table 1:** Antimicrobial Resistance Genes Identified in *P. mirabilis*.

| Sl No. | Resistance Gene | Identity (%) | Phenotype — Antibiotic(s) |
| --- | --- | --- | --- |
| 1 | aph(6)-Id | 100.00% | Streptomycin |
| 2 | aph(3'')-Ib | 100.00% | Streptomycin |
| 3 | bla TEM-1B | 100.00% | Amoxicillin, Ampicillin, Piperacillin, Ticarcillin, Cephalothin |
| 4 | cat | 98.02% | Chloramphenicol |
| 5 | qnrD3 | 100.00% | Ciprofloxacin |
| 6 | sul1 | 100.00% | Sulfamethoxazole |
| 7 | sul2 | 100.00% | Sulfamethoxazole |
| 8 | tet(J) | 99.50% | Tetracycline, Doxycycline |
| 9 | dfrA7 | 100.00% | Trimethoprim |

This dot plot illustrates the specific relationship between each identified resistance gene (y-axis) and its associated antibiotic class (x-axis). Each dot represents a unique gene-class relationship, with colors corresponding to the antibiotic class. This figure provides a clear, one-to-one mapping of the resistance genes to their functional phenotypes, visually summarizing the isolate’s multidrug-resistant profile.

### 3.5. Deciphering Pathogenicity: The Virulence Repertoire of this isolate Defines Its Pathogenic Potential

Further analysis revealed that a broad repertoire of virulence factors are present in the genome, which collectively contribute to its ability to colonize, evade host defenses, and cause disease. These genes are crucial for understanding the pathogen’s transition from a commensal to a clinically significant threat. Genome-wide annotation of the isolate revealed 111 putative virulence-associated genes distributed across multiple functional categories. The largest proportion comprised fimbrial and pili-associated adhesins, including *fim*, *pil*, *pmf*, *sfa*, and autotransporter genes, which are central to surface adhesion and biofilm formation. A complete urease operon (ureA–G) was detected consistent with the strong urease activity that contributes to urinary alkalinization and stone formation, a hallmark of *Proteus* pathogenesis. Genes encoding flagellar components (*fli*, *flg*, *mot*) and chemotaxis proteins were also present, underscoring the swarming motility phenotype of the species. Several toxin and protease-related genes were identified, including *hpmA/B* (hemolysin), *zapA* (metalloprotease), and other protease annotations, supporting roles in host tissue damage and immune evasion. In addition, the genome contained a diverse repertoire of iron acquisition systems, such as TonB–ExbB/ExbD transporters, heme receptors, and siderophore-associated genes. Multiple lipopolysaccharide (LPS) and capsule biosynthesis genes (*wzx*, *wzy*, *waa*, *lpx*, *kps*) were also detected, which may enhance serum resistance and immune evasion.

The analysis further revealed the presence of secretion system genes, including type II, V, and VI system components (*gsp*, *hcp*, *vgrG*, autotransporters), suggesting a versatile secretory capacity for effector proteins. Genes implicated in quorum sensing and biofilm regulation (*luxS*, *pga*, *bcs*) were also identified. Overall, the virulence gene profile (Table 2) highlights a multifactorial pathogenic potential, combining strong adhesive and biofilm-forming ability, robust urease activity, motility, toxin production, iron scavenging, and capsule biosynthesis. These features collectively explain the clinical persistence and pathogenic versatility of this isolate.

**Table 2:** Virulence-associated genes identified in the *P. mirabilis* isolate.

| Sl. No. | Virulence category | Representative genes/loci (from annotation) | Functional role in pathogenesis |
| --- | --- | --- | --- |
| 1 | Adhesins, fimbriae, pili | *fimA*, *fimC*, *sfaA*, *lpfA*, *mrpA*, *pmfA*, autotransporter (*ehaG*) | Adhesion to host cells, biofilm formation, persistence on catheters |
| 2 | Flagella and motility | *fliC*, *flgG*, *flhA*, *motA*, *cheA* | Swarming motility, colonization, dissemination |
| 3 | Urease | *ureA*, *ureB*, *ureC*, *ureE–G* | Urease activity, urinary alkalinization, stone formation |
| 4 | Toxins and proteases | *hpmA*, *hpmB*, *zapA*, metalloprotease genes | Hemolysis, cytotoxicity, tissue damage, immune evasion |
| 5 | Iron acquisition systems | *tonB*, *exbB*, *exbD*, *fecA*, *chuA*, siderophore receptors | Iron uptake under host-limited conditions |
| 6 | LPS and capsule | *wzx*, *wzy*, *waaC*, *lpxA*, *kpsM* | Serum resistance, immune evasion, structural variability |
| 7 | Secretion systems | *gspD*, *gspE* (T2SS), *hcp*, *vgrG* (T6SS), autotransporters (T5SS) | Secretion of adhesins, proteases, effectors |
| 8 | Outer membrane proteins | *ompA, ompC, pal* | Nutrient uptake,<br>immune evasion,<br>adhesion |
| 9 | Quorum sensing & biofilm | *luxS, pgaA–D, bcsA* | Biofilm maturation,<br>population signaling |
| 10 | Stress/serum resistance | *sodB, katG, pmrA/B* | Resistance to oxidative<br>stress and host<br>defenses |

### 3.6. MGEs are Associated that may Drive Horizontal Gene Transfer events

Using IslandViewer4, we identified eight genomic islands in the genome of our *P. mirabilis* isolate that carry clusters of genes associated with both AMR and virulence. We deem that the presence of these islands is significant because they are known hotspots for HGT, serving as reservoirs for novel adaptive traits and confirm the genome’s plasticity, indicating that it can easily acquire new genes from other bacteria. This process, driven by MGEs like genomic islands and insertion sequences, allows a bacterium to rapidly evolve from a harmless commensal to a virulent pathogen. Our findings underscore that HGT is a key mechanism driving the co-evolution of virulence and AMR in this specific *P. mirabilis* isolate (Fig. 4).

**Figure 4.**
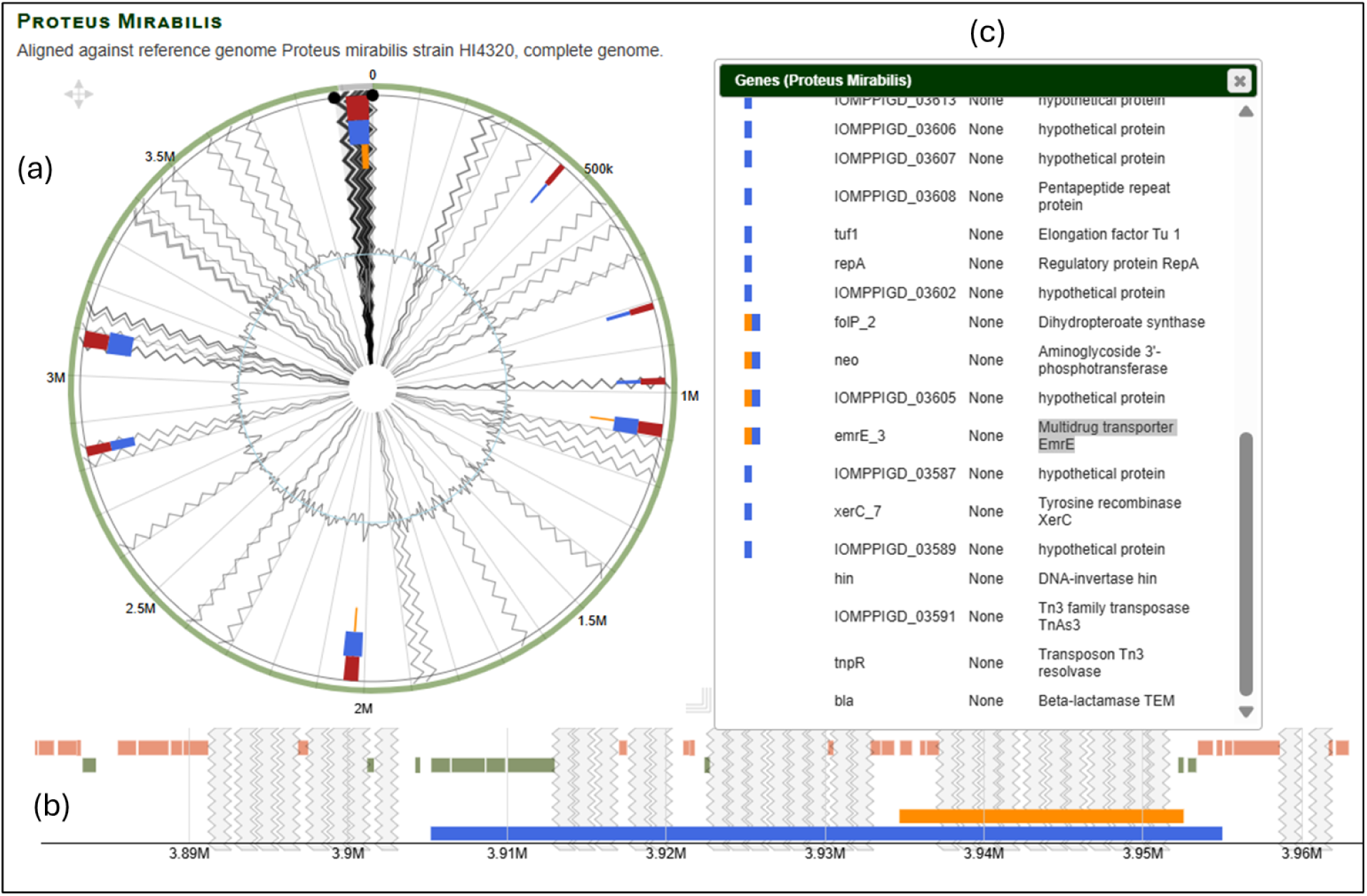
Genomic islands (GIs) predicted in the genome of *P. mirabilis* Indica using IslandViewer (a) Circular representation of the complete genome showing the distribution of predicted genomic islands. GIs are highlighted with colored blocks corresponding to different prediction methods: IslandPick (green), IslandPath-DIMOB (blue), SIGI-HMM (orange), Islander (turquoise), and integrated predictions (dark red). Virulence genes (purple for curated, light purple for homologs), antimicrobial resistance (AMR) genes (pink for curated, light pink for homologs), and pathogen-associated genes (yellow) are visualized as circular glyphs. (b) Horizontal genome view showing zoomed-in regions enriched in predicted GIs, particularly between ∼3.89–3.96 Mb, where multiple clusters of resistance genes and MGEs are observed, suggesting a hotspot for horizontal gene transfer and genomic plasticity. (c) Gene content analysis of predicted GIs, listing several hypothetical proteins along with key resistance and mobility-associated genes, including blaTEM-1B, aminoglycoside resistance determinants, transposases, recombinases, and multidrug transporters (e.g., EmrE). These findings highlight the role of genomic islands in shaping the adaptive evolution of *P. mirabilis* Indica, particularly in enhancing its antimicrobial resistance and virulence potential.

The identification of the IS91 family of insertion sequences in this *P. mirabilis* isolate is particularly noteworthy. Members of the IS91 family are characterized by their rolling-circle (RC) transposition mechanism, which is distinct from the classical “cut-and-paste” or “copy-and-paste” transposition mechanisms employed by most other IS families. This RC-mediated process allows IS91 elements to mobilize adjacent DNA segments, including AMR genes, in a manner analogous to plasmid rolling-circle replication. In this genome, along with other IS elements, IS91 was detected on contig 37, where it is located in close physical proximity to the AMR genes, including blaTEM-1B gene, a β-lactamase conferring resistance to penicillins and related antibiotics. The juxtaposition of an IS91 element near blaTEM-1B has important biological implications. The co-localization of IS91 and blaTEM-1B on contig 37 suggests a potential genetic platform for mobilization and dissemination of β-lactam resistance (Table 3). This highlights the dynamic role of insertion elements in shaping the resistome of *P. mirabilis* and warrants close surveillance, especially considering their contribution to AMR gene spread in healthcare settings.

**Table 3.**
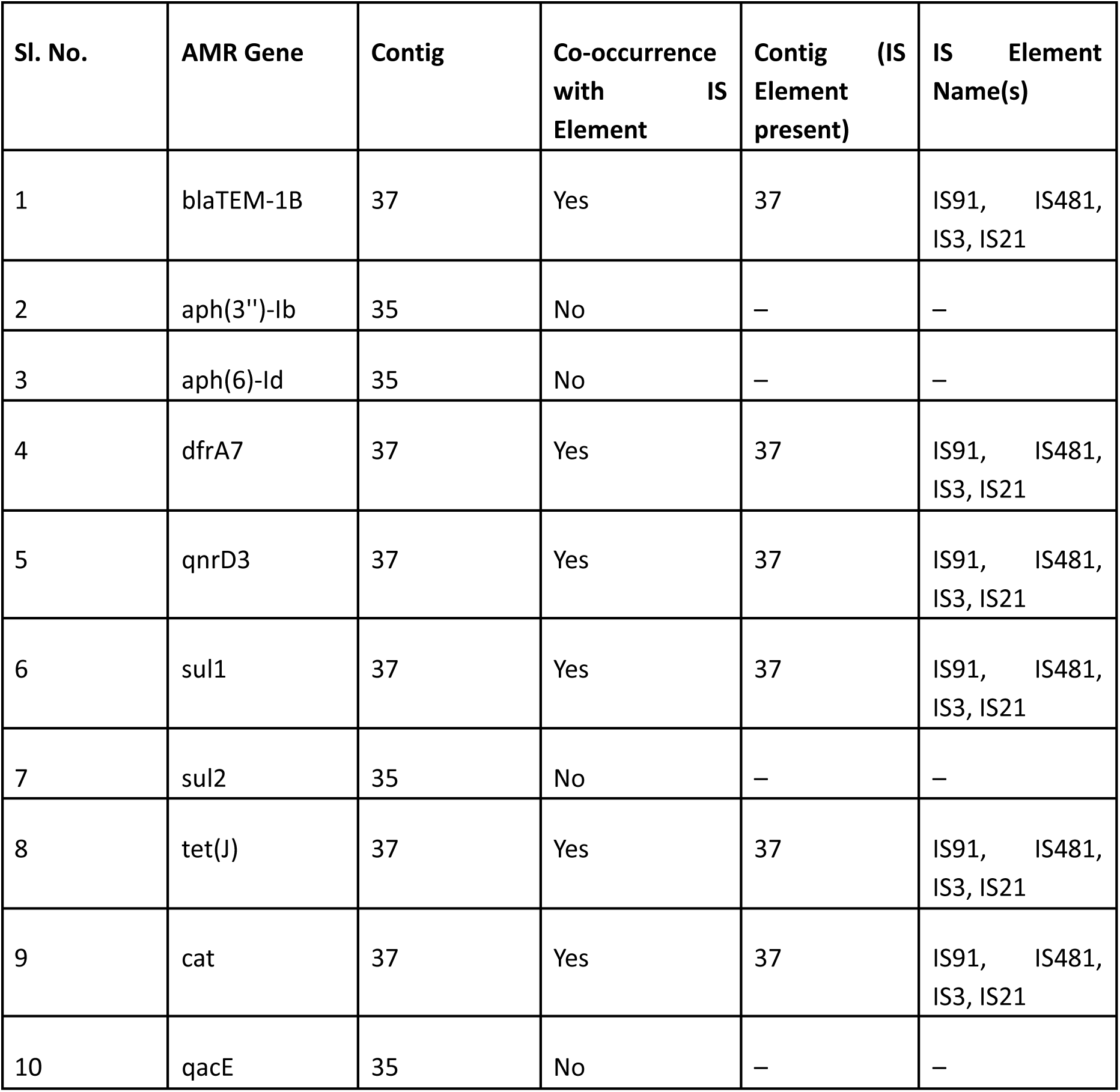
Colocalization of AMR genes with mobile genetic elements (MGEs) in *P. mirabilis* ST-675.

### 3.7. Protein–protein interaction network of *P. mirabilis* Indica showing urease, fimbrial, and hemolysin clusters as key virulence hubs

The STRING interaction network of the *P. mirabilis* Indica (Fig. 5A and 5B) isolate revealed distinct functional clusters of virulence-associated proteins. A urease module (ureA–E) showed dense interconnections, underscoring the importance of urease in urinary tract colonization and stone formation. A fimbrial operon cluster (mrpA–H) formed another well-defined module, reflecting its role in adhesion and biofilm formation. Hemolysin genes (hpmA and hpmB) emerged as central hubs with extensive links to both metabolic and virulence proteins, suggesting coordinated functions in host damage, nutrient acquisition, and immune evasion. Peripheral proteins, including zapA, were less connected but indicated specialized roles in tissue invasion. Collectively, the network highlights a modular but interconnected virulence architecture that equips the isolate for persistence in hostile host environments.

**Figure 5.**
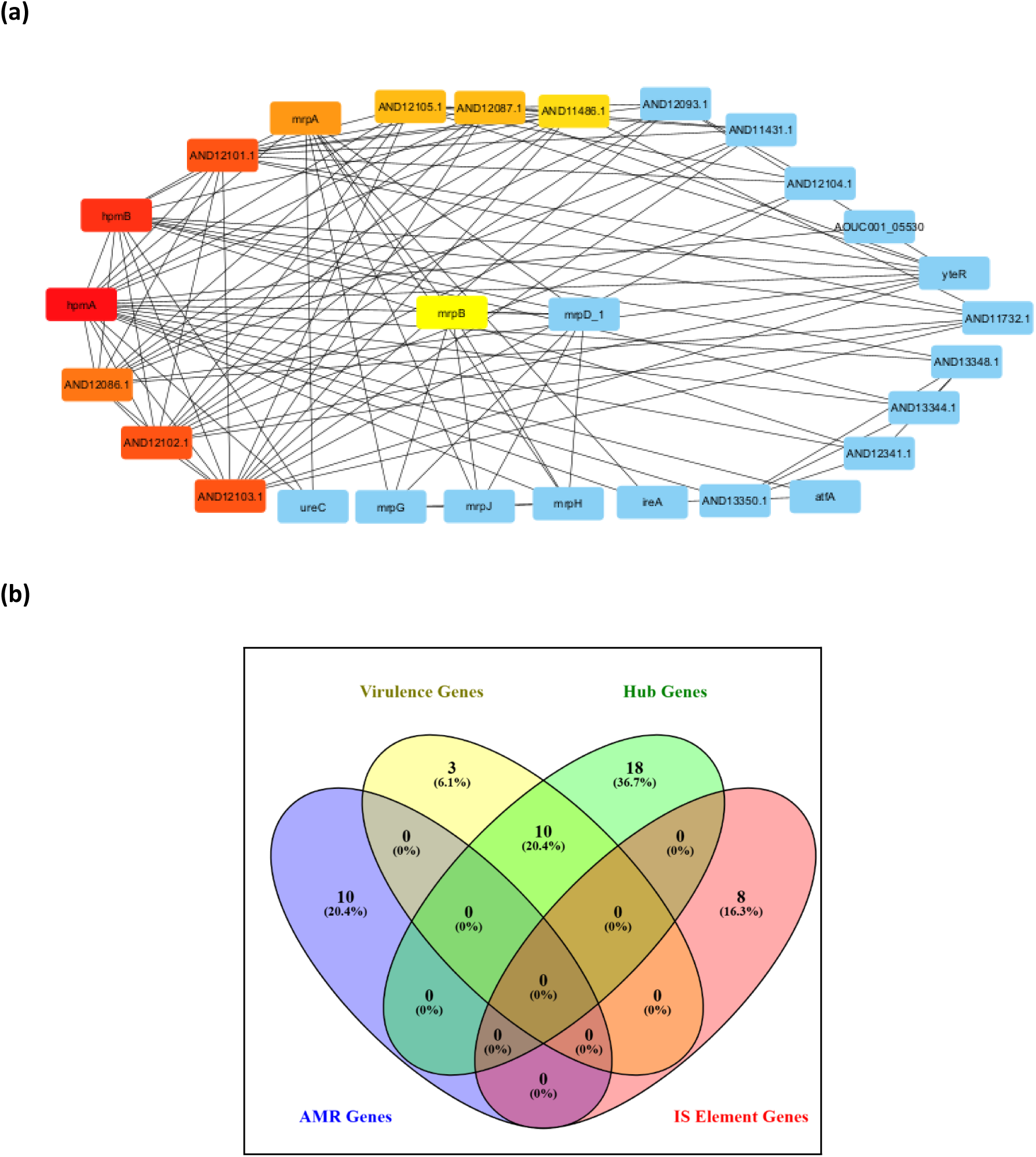
(a) Protein–protein interaction (PPI) network of *P. mirabilis* virulence-associated genes constructed using STRING and analyzed with CytoHubba. Node color intensity corresponds to interaction importance, with darker red/orange indicating higher centrality. Other associated genes include urease components (ureC), fimbrial subunits (mrpG, mrpJ, mrpH, mrpD), and adhesion-related proteins (ireA, atfA, yteR), shown as blue nodes, which contribute to colonization and persistence. Yellow nodes represent additional hub candidates identified by network centrality. (b) Comparative Venn diagram of hub genes, antimicrobial resistance (AMR) genes, virulence factors, and insertion sequence (IS) elements in *P. mirabilis* isolate.

The interaction network highlights the central role of the hemolysin genes hpmA and hpmB, together with the fimbrial operon regulator mrpA, as major hubs (red–orange nodes) strongly connected to multiple partners indicating that these are key mediators of host–pathogen interactions, inflammation, and tissue damage during infection. underscoring the convergence of fimbrial and hemolysin pathways in shaping *P. mirabilis* pathogenicity, particularly in urinary tract infections and prostatitis. The Venn diagram illustrates the overlap and unique distribution of hub genes (identified by CytoHubba analysis), AMR genes (detected by ResFinder), virulence-associated genes (from VFDB), and IS elements (from ISFinder). Notably, hub genes showed the highest intersection with virulence-associated genes, underscoring their dual contribution to both pathogenic potential and network centrality. Nevertheless, limited overlap was observed with AMR genes and IS elements, indicating distinct functional roles with occasional convergence in bacterial adaptation and survival.

### 3.8. Comparative Genomics and Phylogenetic Analysis clustered this isolate firmly within the Proteus clade

Core-genome-based phylogenetic analysis confirmed the evolutionary placement of our isolate. As shown in Figure 6, our isolate, designated *P. mirabilis* Indica, clustered firmly within the *Proteus* clade, distinct from other genera like *Providencia* and *Xenorhabdus*. The short branch length leading to our isolate indicates its close genetic relationship with other *P. mirabilis* strains. This analysis provides a visual confirmation of the isolate’s species identity and validates its unique designation as a subspecies potentially endemic to the Indian subcontinent.

**Figure 6.**
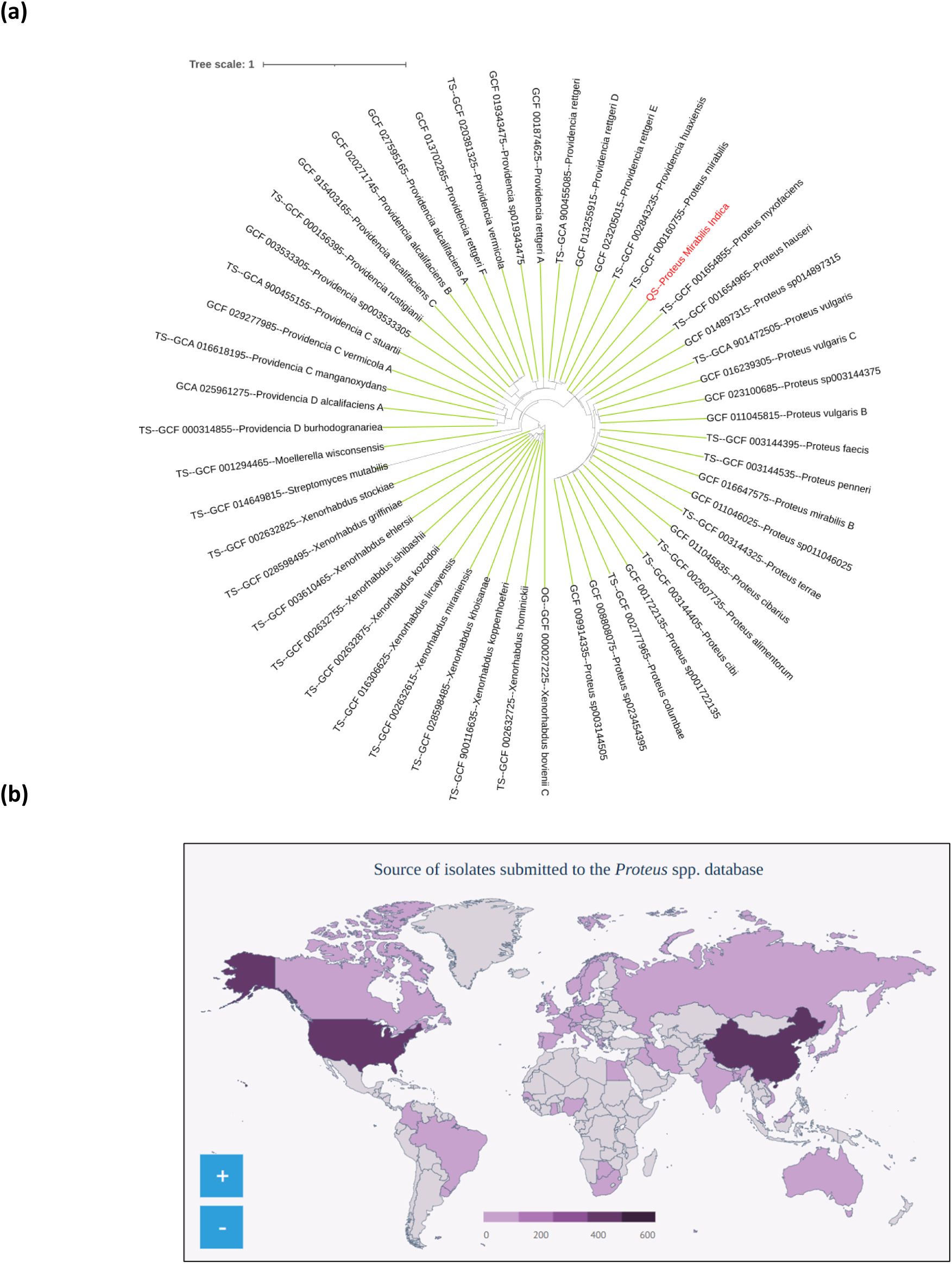
(a) Core-Genome Phylogenetic Tree of the *P. mirabilis* Isolate. (b) Global distribution of *Proteus* spp. isolates submitted to the PubMLST database.

A core-genome-based Maximum Likelihood phylogenetic tree illustrating the evolutionary relationship of our isolate (labeled in red) with other reference strains. The tree is rooted with an outgroup of *Providencia* and *Xenorhabdus* species. The branch lengths represent the number of nucleotide substitutions per site, as indicated by the scale bar (0.05). The clustering of our isolate within the *Proteus* clade confirms its species identity. Further, the geographic distribution of *Proteus* spp. isolates submitted to a global database highlight regional differences in reporting and surveillance (Fig. 6b). The shading intensity reflects the number of isolates, with darker colors indicating higher submissions with the United States and China standing out as the largest contributors, each with more than 600 isolates, followed by several European and Asian countries with moderate submissions. In contrast, large regions of Africa, the Middle East and parts of South America report relatively few isolates, suggesting either a genuinely lower detection rate or, more likely, underrepresentation due to limited genomic surveillance and reporting infrastructure.

## Discussion

### Virulence determinants were distinct in the *P. mirabilis Indica*

This study presents the first genomic characterization of a *P. mirabilis* ST-675 isolate harboring *blaTEM-1B* from India, providing important insights into the convergence of AMR and virulence in this opportunistic pathogen. The combination of WGS, phylogenetic analysis and virulome–resistome profiling underscores how *P. mirabilis* continues to adapt within clinical environments through HGT and the acquisition of MGEs. These findings extend our understanding of the evolutionary trajectory of *P. mirabilis* in India, a region where genomic data for this species remain limited despite its recognized role in urinary tract and bloodstream infections. What we perceived from our study is that the resistome of the isolate revealed a broad repertoire of AMR genes, most notably *blaTEM-1B*, a β-lactamase widely distributed among Enterobacterales. While carbapenemases were absent, the co-localization of resistance determinants with insertion sequences (IS elements) on Contig 37 provides direct evidence of genomic mobility, suggesting a mechanism for the coordinated dissemination of both resistance and virulence traits. This observation is consistent with recent reports where IS elements facilitate the mobilization of AMR loci, enabling rapid adaptation to antibiotic pressure (9,14). On the other hand, comparisons with the HI4320 reference strain illustrate this evolutionary expansion. HI4320, isolated in 1982 from the urine of a long-term catheterized elderly woman in Baltimore, Maryland, carried only *cat* (M11587) and *tet(J)* (ACLE01000065) as its principal resistance determinants (29,30). By contrast, the present Indian isolate (“*P. mirabilis Indica*,” 2025) displays a MDR phenotype, likely the outcome of cumulative gene acquisition driven by antibiotic exposure in hospital settings. This transition reflects a broader epidemiological trend where originally susceptible strains evolve into MDR clones, complicating treatment regimens and infection control.

The virulome of *P. mirabilis Indica* was equally notable. The identification of multiple fimbrial adhesins indicates a strong capacity for adherence and biofilm formation, critical for CAUTIs and persistent colonization (1). The presence of *ail* genes, traditionally linked to invasive phenotypes in enteric pathogens such as *Yersinia*, points to an expanded invasive potential that is increasingly recognized in clinical *P. mirabilis* isolates. Hemolysin and other toxin-encoding genes further underscore the destructive capacity of this strain, which can damage host tissue and evade immune clearance. Whereas the MLST and genotyping of P mirabilis yielded distinct pathways, what we sought to ask was whether or not any genes, by and large, have interactions with pathways other than mere associations corroborating the hypothesis that the top AMR/MDR genes are largely indispensable for infection. Therefore, our integrated systems biology approaches using STRING, Cytoscape-Cytohubba highlighted the functional integration of virulence determinants in the *P. mirabilis* Indica isolate. We identified Urease and fimbrial clusters that reflected its ability to colonize the urinary tract, form biofilms, and promote stone development, while hemolysin genes emerged as central hubs bridging metabolic and pathogenic pathways. Peripheral factors such as zapA further contributed to tissue damage, indicating specialized but complementary roles. This coordinated network not only explains the bacterium’s persistence in immunocompromised hosts but also pinpoints key nodes such as urease, fimbrial adhesins, and hemolysins that may serve as promising targets for vaccine design or novel therapeutics. The central positioning of mrpA, hpmA and hpmB as key interactors highlights their critical contribution to *P. mirabilis* pathogenesis. It may be possible that as the mrpA gene encodes the major structural subunit of mannose-resistant Proteus-like fimbriae, it mediates adhesion to uroepithelial and prostate epithelial cells, thereby facilitating colonization and persistence within the urinary tract. On the other hand, hpmA encodes a hemolysin and hpmB functions as its activator/translocator; and together, these genes contribute to cytotoxicity by inducing pore formation in host cell membranes, triggering inflammation, and promoting tissue damage. Their combined activity not only possibly enhances bacterial survival by evading host defenses but also exacerbates inflammatory responses within the prostate, a hallmark of chronic bacterial prostatitis. The dual role of these genes in both adhesion and cytotoxicity underscores their importance in host–pathogen interactions and identifies them as pivotal virulence determinants driving *P. mirabilis*–associated urinary tract infections and prostatitis. Taken together, these virulence features create a multifactorial arsenal that synergizes with AMR determinants. For example, biofilm formation not only protects bacteria from host immunity but also reduces antibiotic penetration, amplifying the challenge of eradicating infections. The interplay between virulence and resistance therefore positions *P. mirabilis* as both a therapeutic challenge and a persistent nosocomial pathogen.

### Resistome of the isolate revealed a broad repertoire of AMR genes

The identification of eight genomic islands in the Indian isolate provides further evidence of genomic plasticity and ongoing adaptive evolution. These islands contain a mix of hypothetical proteins, AMR determinants, and mobility-associated genes, reinforcing the role of HGT in shaping bacterial fitness. Notably, the clustering of genomic islands around specific chromosomal regions highlights potential hotspots for recombination and gene acquisition, echoing patterns observed in other Gram-negative pathogens. Phylogenetic comparisons revealed relatedness to a Japanese clinical isolate, suggesting transnational or pan-Asian connectivity of high-risk *P. mirabilis* clones. This finding is consistent with the globalization of multidrug-resistant Enterobacterales, where international travel, medical tourism, and shared antibiotic practices foster the spread of resistant lineages across borders. The clinical implications of these findings are profound, particularly for immunocompromised and cancer patients. Such patients are highly susceptible to opportunistic infections, and impaired immunity reduces their ability to contain bacterial invasion. The coexistence of multiple AMR genes with a broad virulence repertoire therefore increases the likelihood of persistent colonization, recurrent infections, and poor clinical outcomes. Of particular concern is the detection of *qacE* efflux genes, which confer tolerance to quaternary ammonium compounds. These disinfectants are widely used in hospital settings, including oncology and transplant wards. Resistance to disinfectants undermines standard infection-prevention measures and heightens the risk of nosocomial transmission in high-risk units (4). The study by Sharon et al. (32) provides additional context. They reported a high prevalence of pro-inflammatory uropathogens, particularly *Escherichia coli*, in men with prostate cancer (PCa), supporting the association between infection, chronic inflammation, and cancer progression. Although *P. mirabilis* was detected at a lower frequency (2.8%), its biofilm-forming ability, urease production, and multidrug resistance render it a clinically important pathogen even at low prevalence. In the context of PCa and immunosuppression, such traits can aggravate urinary tract infections, perpetuate inflammation, and complicate therapy. Thus, the presence of *P. mirabilis* in these patients warrants greater clinical attention than prevalence data alone may suggest.

### Taxonomic classification suggests a divergent sub-variant possibly leading to HGTed events

Taxonomic classification confirmed the isolate as *P. mirabilis* with 100% confidence, firmly establishing its identity as an opportunistic uropathogen. The MLST assigned it to ST-675, with close relatedness to ST-797, suggesting a divergent sub-variant by microevolution and HGT. Virulence typing (vST-1157) revealed a distinct profile, characterized by virulence determinants not typically seen in reference ST-675 isolates. This divergence highlights the dynamic nature of *P. mirabilis* evolution, where microevolutionary processes continually drive the shift from less pathogenic to more virulent strains. The co-occurrence of AMR and virulence traits in this isolate reflects a broader trend across Enterobacterales, where resistance and pathogenicity increasingly converge in high-risk clonal variants.

Noncoding RNAs are increasingly recognized as fine-tuners of bacterial virulence and metabolic adaptation, often responding to environmental cues such as pH, nitrogen availability, and host immune stress. Such an ncRNA could represent a novel layer of post-transcriptional regulation of urease activity, impacting P. mirabilis pathogenesis in urinary tract infections and possibly influencing persistence in the host. (39,40). In this isolate several regulatory RNAs are present among them GlmZ and GlmY play important roles in AMR. They might modulate peptidoglycan biosynthesis by activating *glmS*, thereby influencing susceptibility to β-lactam antibiotics. Similarly, GcvB RNA regulates amino acid transporters and outer membrane proteins, which might be linked to efflux and permeability changes contributing to multidrug resistance. The putative aminoglycoside riboswitch/attI site is particularly noteworthy, as it may regulate integron-associated resistance cassettes, highlighting a direct link between riboswitch activity and acquisition of AMR genes (Supplementary Data 1). Stress-responsive RNAs such as RyhB, RybB, Spot42, and cspA thermoregulator also contribute indirectly by modulating iron metabolism, oxidative stress, and membrane porin expression, thereby facilitating bacterial survival under antibiotic pressure. Additionally, the Guanidine-II riboswitch has been associated with efflux pump regulation, further strengthening the role of riboswitches in drug tolerance. Together, these findings underscore that lncRNAs in *P. mirabilis* are not merely passive elements but they might be active regulators of host–pathogen interactions and antibiotic stress responses, and may represent novel targets for therapeutic intervention.

In contig 10, we identified a noncoding RNA associated with the *ure* operon, which might be associated with the expression of urease, a key virulence factor in *P. mirabilis*. Its genomic proximity suggests a possible regulatory role in modulating urease expression in response to host conditions. Given that bacterial ncRNAs often fine-tune virulence and stress adaptation, this ncRNA may represent a novel post-transcriptional regulator influencing *P. mirabilis* persistence in urinary tract infections. Functional studies are needed to confirm its role.

### Future perspectives and limitations

From a public health perspective, the emergence of multidrug-resistant *P. mirabilis* lineages with expanded virulence capacity raises concerns for global dissemination and clinical management. Integrating genomic surveillance with antimicrobial stewardship is essential to anticipate these threats. Region-specific monitoring in India is particularly critical, given the country’s high antibiotic consumption, dense hospital environments, and rising burden of MDR infections. Beyond surveillance, WGS datasets provide a foundation for translational applications. Immunoinformatics-based approaches can leverage genomic data to identify vaccine candidates, while phage therapy and monoclonal antibodies offer promising alternatives to traditional antibiotics. At the same time, stewardship programs must carefully balance the use of existing antibiotics to slow the selection pressure driving resistance. However, our work is void of a few limitations, *viz.*. (a) First, the analysis was based on a limited sample size, which may restrict the generalizability of the findings to broader *P. mirabilis* populations or other clinically relevant pathogens. Second, while genomic and regulatory RNA features were characterized, detailed host–pathogen interaction studies were not performed, leaving gaps in understanding how these regulatory elements influence bacterial survival and immune evasion.

## Conclusions

This study presents the first genomic characterization of a *P. mirabilis* ST-675 isolate carrying *blaTEM-1B* from India, providing evidence of the convergence of antimicrobial resistance, virulence, and mobile genetic elements within a single high-risk lineage. Comparative analysis highlights a striking expansion of the resistome beyond the historical HI4320 reference strain, alongside a virulence repertoire enriched with fimbrial adhesins, toxins, and invasion-associated genes. The identification of multiple genomic islands and the co-localization of AMR genes with insertion sequences underscore the role of horizontal gene transfer and microevolution in shaping this pathogen’s adaptability.

Clinically, the coexistence of multidrug resistance, virulence factors, and disinfectant tolerance mechanisms renders this isolate particularly concerning in hospital settings, where it threatens to persist despite therapy and infection-control measures. The risks are magnified for cancer patients and immunocompromised hosts, in whom impaired immunity provides a foothold for opportunistic pathogens with both resistance and virulence attributes. Beyond posing an immediate therapeutic challenge, this strain also represents a potential reservoir for the dissemination of clinically relevant genes across bacterial populations.

Taken together, these findings underscore the urgent need for region-specific and global genomic surveillance of P. mirabilis to better track emerging high-risk clones, inform infection control measures, and safeguard vulnerable patient populations such as cancer and immunocompromised hosts. Integration of whole-genome sequencing with antimicrobial stewardship will be critical to anticipate emerging high-risk clones and to inform clinical management. At the same time, the genomic insights gained here provide a foundation for the exploration of alternative strategies, including vaccine development, bacteriophage therapy, and immunotherapeutics, to counter the growing threat of multidrug-resistant *P. mirabilis* in vulnerable patient populations.

## Data Availability

Data has been submitted to NCBI-SRA. The raw reads of sequence data could be accessed from SRA through BioProject ID: PRJNA1206681. SRA records will be accessible with the following link after the indicated release date: https://www.ncbi.nlm.nih.gov/sra/PRJNA1206681.

## Authors Contribution

SA, PK, PS, and GC contributed to the study design. LLA, BC, PK, and SA collected samples. Data were analyzed and interpreted by SA, PK, SD1, PS, SB and GC. SA and PK designed the figures. The manuscript was written by SA, PK, and edited by SD1, MR, SD2, PS, and GC. GC and PS supervised the study. All authors had full access to all the data in the study and accepted responsibility for the decision to submit for publication. SA & PK contributed equally to this study.

## Ethics statement

This study was approved by the Institutional Ethics Committee of IQ City Medical College & NH Hospital, Durgapur, India (Approval No. IQMC/IEC/Project/17/28) for the genomic characterization of *Acinetobacter baumannii*. During subsequent laboratory analysis, the isolate was found, through whole-genome sequencing, to be *P. mirabilis* rather than *A. baumannii*. The sequencing data and downstream analyses presented in this manuscript, therefore, pertain to *P. mirabilis*. Isolate in this study was recovered from discarded clinical culture plates without access to any patient-identifiable information. According to the International Ethical Guidelines for Health-related Research Involving Humans (CIOMS-WHO, 2016) and the Indian Council of Medical Research (ICMR) guidelines, such use of anonymized bacterial isolates without any associated personal data does not require individual patient consent, as it does not constitute human subject research.

In accordance with the International Ethical Guidelines for Health-related Research Involving Humans (CIOMS-WHO, 2016) and the Indian Council of Medical Research (ICMR) guidelines, the use of anonymized bacterial isolates without linked personal data does not constitute human subject research and therefore does not require individual informed consent. We have carefully followed the

1. CIOMS-WHO (2016) International Ethical Guidelines for Health-related Research Involving Humans https://cioms.ch/publications/

➤ Guideline 3: Use of de-identified, minimal-risk samples may not require consent.
2. ICMR National Ethical Guidelines (2017) https://ethics.ncdirindia.org/

➤ Section: “Biological Materials and Data” permits use of unlinked, anonymized clinical samples without consent under specified conditions.
3. U.S. DHHS – Common Rule (45 CFR 46) (For international reviewers) https://www.ecfr.gov/current/

➤ Research on non-identifiable biospecimens is not considered human subject research.

